# Visualizing the periodic Ribo-seq reads with RiboPlotR

**DOI:** 10.1101/694646

**Authors:** Hsin-Yen Larry Wu, Polly Yingshan Hsu

**Affiliations:** Department of Biochemistry & Molecular Biology, Michigan State University, East Lansing, MI 48824 USA

## Abstract

**Background:** Ribo-seq has revolutionized the study of mRNA translation in a genome-wide scale. High-quality Ribo-seq data display strong 3-nucleotide (nt) periodicity, which corresponds to translating ribosomes decipher three nucleotides each time. While the 3-nt periodicity has been widely used to study novel translation events and identify small open reading frames on presumed non-coding RNAs, tools which allow the visualization of those events remain underdeveloped.

**Findings:** RiboPlotR is a visualization package written in R that presents both RNA-seq coverage and Ribo-seq reads for all annotated transcript isoforms in a context of a given gene. In particular, RiboPlotR plots Ribo-seq reads mapped in three reading frames using three colors for one isoform model at a time. Moreover, RiboPlotR shows Ribo-seq reads on upstream ORFs, 5’ and 3’ untranslated regions and introns, which is critical for observing new translation events and potential regulatory mechanisms.

**Conclusions:** RiboPlotR is freely available (https://github.com/hsinyenwu/RiboPlotR) and allows the visualization of the translating features in Ribo-seq data.

## INTRODUCTION

mRNA translation is the last step of the central dogma of molecular biology. Despite its importance in directly control protein production, it is much less understood compared to transcription. Ribosome profiling, also known as Ribo-seq, has revolutionized the study of translation by mapping and quantifying ribosome footprints on each mRNA in the past decade [1]. High-quality Ribo-seq data display strong three-nucleotide (nt) periodicity, which corresponds to ribosomes decode three nucleotides per codon. While many computation tools use 3-nt periodicity to facilitate the identification of new translation events in the transcriptome, the visualization of the periodicity of Ribo-seq reads remains difficult. So far, the visualization plots shown in most literature either completely exclude consideration of periodicity, or only focus on periodicity in the context of one mature transcript isoform. In the latter case, the Ribo-seq reads in-frame to each of the three frames of ORFs are shown (**Figure 1A**) [2]. This “single transcript” style of plotting is useful to identify overlapping translation events within a specific mature isoform but would miss Ribo-seq reads that belong to other isoforms, and those mapped in the introns and unannotated coding exons.

**Figure 1.**
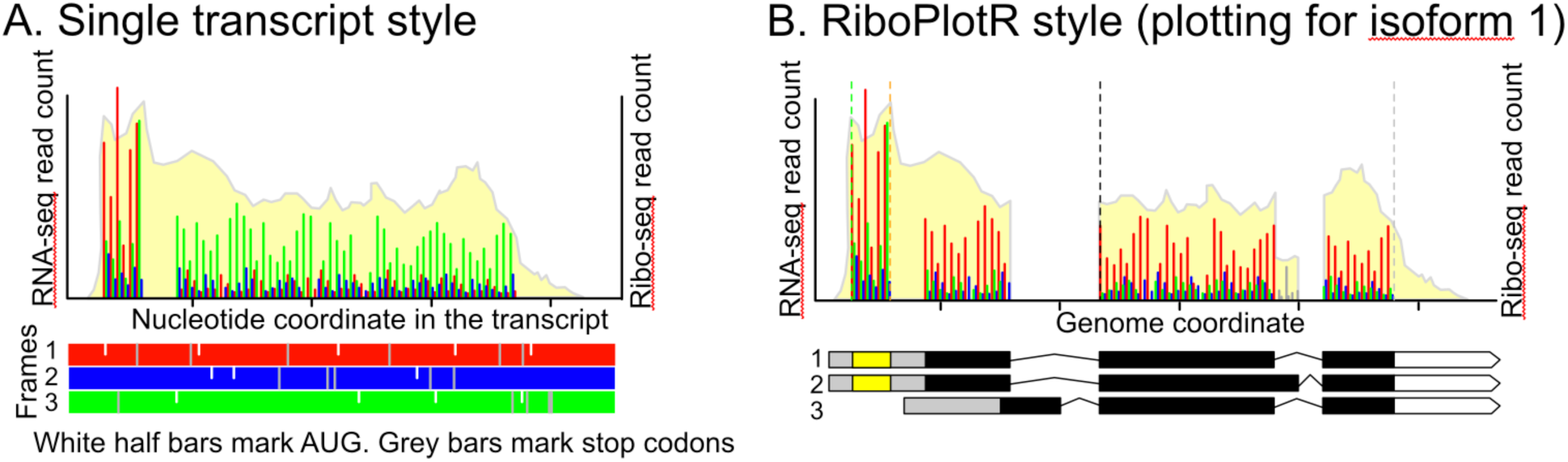
The commonly used single transcript style versus RiboPlotR style. A. The commonly-used single transcript style plot. RNA-seq and Ribo-seq reads are shown for one mature transcript isoform per plot. For Ribo-seq reads, either the most 5’ or the P-site position (the first nt within the peptidyl site within the ribosome) is used for plotting. The first nt of the transcript is considered as frame 1. Thus, the Ribo-seq reads for the annotated CDS could be in one of the three colors. AUG start codons are marked as the white half lines in all three frames. The stop codons are marked as the grey lines in all three frames. B. The RiboPlotR style shows all annotated transcript isoform models in parallel with the RNA-seq coverage and Ribo-seq P-site reads. Within the gene model, the grey boxes indicate 5’ UTR, the black boxes indicate the annotated ORF, and the white pentagon arrow indicates 3’ UTR. In addition to the annotated ORF, one upstream ORF (yellow box in the gene model) could be shown in the same plot. For all transcript isoforms of a given gene, the same RNA-seq coverage and Ribo-seq P-sites are used for plotting. For the annotated ORF, the expected CDS range is marked between a black dashed line (translation start site) and a grey dashed line (translation stop site); while for the uORF, the CDS range is marked between a green dashed line (translation start site) and an orange dashed line (translation stop site). The Ribo-seq P-sites that are mapped in the expected frame, +1 frame, and +2 frame are marked with red, blue, and green, respectively. The Ribo-seq P-sites mapped outside of the expected CDS in either the annotated ORF or uORF are shown in grey. Here the isoform 1 and 2 are both expressed and translated, but the isoform 3 is not expressed. Since we select the isoform 1 for plotting here, some Ribo-seq P-sites that are unique to the isoform 2 and not covered by the isoform 1 are marked in grey.

Here we developed RiboPlotR (**Figure 1B**) for visualizing RNA-seq/Ribo-seq reads in the context of a given gene, including Ribo-seq reads mapped in the intronic and 5’ and 3’ untranslated regions (UTRs), with all transcript isoform models displayed in parallel in the plot. There are several advantages of RiboPlotR style: (1) We can detect novel translation events occurred in the unannotated coding regions, such as those in the introns and UTRs. (2) By including all transcript isoform models in the plot, we can visually determine which transcript isoform(s) is translated in most cases. (3) Comparing the sequencing data and the annotated gene models in parallel, we can easily identify a discrepancy between the Ribo-seq data and the predicted coding sequences (CDSs), such as frameshift and variation in coding regions; similarly, any deviation in the mRNA profile from the annotated transcript isoforms will also be easily visualized. (4) Relative Ribo-seq abundance in different transcript features, such as introns or uORFs, which suggest potential regulatory mechanisms, could be visualized. Below, we describe usages and examples using RiboPlotR to visualize translation events considering a predicted uORF and different transcript isoforms in the context of a gene.

## METHODS

RiboPlotR uses the base R commands for plotting. It requires the *GenomicRanges, GenomicFeatures, GenomicAlignments, Rsamtools* and *rtracklayer* packages from Bioconductor [3–5]. RiboPlotR reads in the transcriptome annotation GFF or GTF files with gene.structure function, which is based on the makeTxDbFromGFF function from GenomicFeatures. The gene.structurefunction further processes the transcriptome database to obtain the genomic coordinates of transcripts, exons, CDSs, and UTRs that belong to each gene. To visualize the uORFs, another GFF/GTF could be load with the uorf.structure function. In the current version of RiboPlotR, users could visualize one uORF at a time. The rna_bam.ribo function reads in the RNA-seq mapped bam file(s) and the Ribo-seq P-site coordinate file(s). The Ribo-seq P-site is denoted as the first nt of the peptidyl-site within the ribosome. The RNA-seq bam files need to be mapped and sorted by genome coordinates. A *bai* (i.e., bam index) file is also required. We recommend the users map the RNA-seq reads with the *STAR* aligner and sort/index the bam file with *samtools* [6, 7].

The Ribo-seq P-site coordinate file should be a tab-delimited file. From left to right, the Ribo-seq P-site coordinate file should contain “count”, “chromosome number”, “coordinate” and “strand” information but without a header. For each chromosome coordinate, the column “count” should be the sum of Ribo-seq P-site counts. The P-site could be acquired from the RiboTaper output P_sites_all file or any other package that defines the P-site positions. If the P-site information is obtained from RiboTaper, the users could use the following linux command cut -f 1,3,6 P_sites_all | sort | uniq -c | sed -r ‘s/^(*[^]+) +/\1\t/’ > name_output_file to produce the P-site file.

Together, RiboPlotR requires the following input files: (1) a gtf or gff3 for transcriptome annotation, which should be recognizable with the GenomicFeatures package; (2) a mapped and coordinate-sorted bam file(s) for RNA-seq; (3) a tab-delimited file(s) for Ribo-seq P-site coordinate file. Finally, a gtf or gff3 for uORF coordinates is optional. Moreover, users could read in one or two set of the bam and P-site files to compare translation in two different conditions.

RiboPlotR has exported four plotting functions: PLOTc, PLOTt, PLOTc2 and PLOTt2. PLOTc plots RNA-seq and Ribo-seq in one panel (plot **<u>c</u>**ompact). PLOTt plots RNA-seq and Ribo-seq in two panels (plot **t**wo). PLOTc2 plots RNA-seq and Ribo-seq in one panel for two conditions. PLOTt2 plots RNA-seq and Ribo-seq separately for two treatments or conditions.

## RESULTS AND DISCUSSION

### Basic utilities

RiboPlotR could be used to plot the data from different organisms because it uses the base R function to input the Ribo-seq P-site coordinates and the widely-used Bioconductor packages to input standard gtf/gff3 files and RNA-seq bam files. The RiboPlotR package contains a sample Ribo-seq dataset from Arabidopsis root and shoot described in Hsu *et al*. 2016 [8]. The sample RNA-seq data used here is a paired-end 100 nt dataset from the same study but not yet published. We provided two examples below to showcase RiboPlotR.

The basic workflow of RiboPlotR is:

1. Run gene.structure(); load the transcriptome annotation gtf/gff3 file containing the gene, mRNA/transcript, exon and CDS ranges.
2. (Optional) To plot a uORF for a transcript, users will also load the uORF gtf/gff3 file using the uorf.structure() function.
3. Run rna_bam.ribo(); load the mapped and coordinate-sorted RNA-seq bam file and the ribo-seq P-site position file.
4. Use one of the four functions below, enter gene name and isoform number to plot the translation of the isoform.

#### Four styles of the plots are available

PLOTc: plots RNA-seq and Ribo-seq in one panel (plot **<u>c</u>**ompact)

PLOTt: plots RNA-seq and Ribo-seq separately in two panels (plot **t**wo)

PLOTc2: plots RNA-seq and Ribo-seq in one panel for two conditions

PLOTt2: plots RNA-seq and Ribo-seq separately for two conditions

### Plot presentations

RiboPlotR plots each isoform of a given gene separately. Only one isoform would be plotted each time, and the default is to plot the isoform 1. For each isoform, the same RNA-seq and Ribo-seq reads are used for plotting; the only difference is the expected coding region for the Ribo-seq reads, which is indicated by a black dashed line (expected translation start) and a grey dashed line (expected translation stop). Inside the expected coding region, Ribo-seq P-sites that are mapped in the expected frame, +1 frame, and +2 frame are presented in red, blue and green lines, respectively. Ribo-seq P-sites that are outside of the expected coding region are shown in grey. The x-axis below the gene models indicates the genomic coordinates, while the y-axis indicates the Ribo-seq P-site counts. When an isoform is translated, the majority of P-sites should cover the expected coding sequences and are shown in red color. If two isoforms cover a different coding region at the 3’ ends, the two plots will have different coloring scheme at the 3’ end. This design allows users to quickly spot if a plotted isoform is being actively translated (see example below). Below we show two examples to demonstrate how to interpret the RiboPlotR plots.

### Examples of plotting without or with an uORF

Figure 2 plots *AT3G02470*, which encodes the S-AdenosylMethionine DeCarboxylase (SAMDC), using PLOTc function. In addition to the abundant Ribo-seq reads in the expected coding sequence, many reads are present in the 5’ UTR (**Figure 2A**). The Ribo-seq reads in the 5’ UTR imply two possibilities: the 5’ UTR reads could be from an uORF(s) or resulting from the usage of a non-AUG translation start site. *SAMDC* is known to have a <u>c</u>onserved <u>p</u>eptide <u>uORF</u> (CPuORF9), which is annotated as *AT3G02468* in the TAIR10 annotation. We plotted the *AT3G02470* again with the ‘uORF=“AT3G02468”’ option. It is clear that the Ribo-seq reads in the 5’ UTR are highly enriched in the predicted coding sequence for *CPuORF9* and are consistent with the expected reading frame (**Figure 2B**). Thus, *CPuORF9* is highly translated in the condition examined here. This example demonstrates that RiboPlotR is useful for visualizing translation events occurring within a gene in addition to the annotated coding sequence.

**Figure 2.**
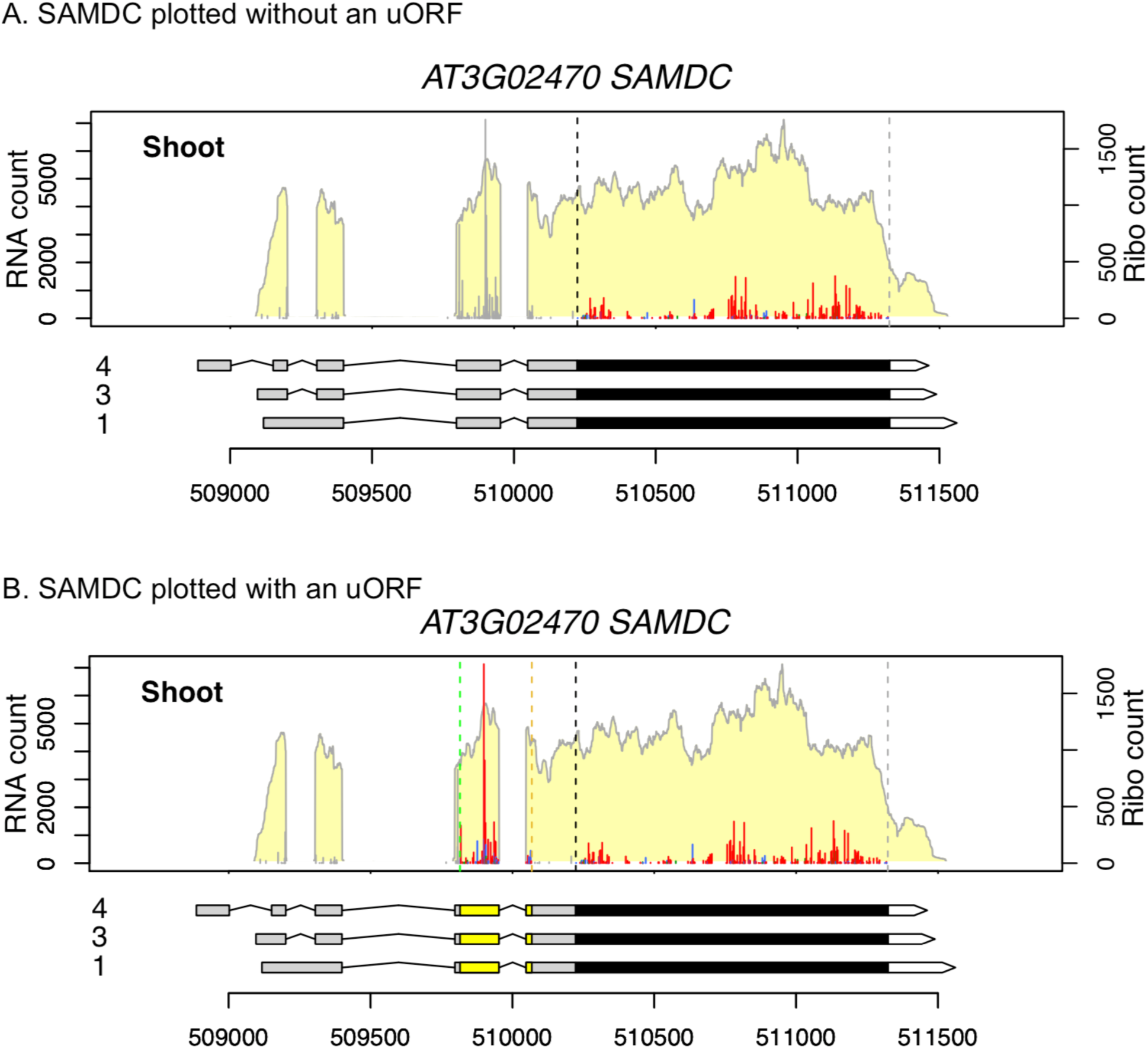
An example of plotting without or with an uORF. A. The *AT3G02470* gene is plotted using PLOTc function with RiboPlotR. Only the annotated ORF is considered. The Ribo-seq P-site reads outside of annotated CDS are marked as grey. B. Plot both the annotated ORF and the CPuORF9 with RiboPlotR. Data presentation are the same as described in the Figure 1 legend; genomic coordinates are shown below the gene models.

Some transcript isoforms have an identical CDS, and their only differences are located in the UTRs. In these cases, we could use the RNA-seq plot to determine which isoform(s) is expressed. For example, the three isoforms for *SAMDC* share the same CDS (**Figure 2**). However, based on the RNA-seq coverage around the exon-intron structures, it is clear only the isoform 3 is expressed.

### Examples of plotting different transcript isoforms

Figure 3 plots *AT4G21910*, which encodes a MATE (the <u>M</u>ultidrug <u>A</u>nd <u>T</u>oxic compound <u>E</u>xtrusion) efflux family transporter, using PLOTc2 function comparing data in the shoot and root. In this case, the first two isoforms are the major translated isoforms since the RNA-seq/Ribo-seq coverage and frame color agree with these two isoforms (**Figure 3A and 3B**). Moreover, the isoform 1 is preferentially transcribed and translated in the shoot, while the isoform 2 is preferentially transcribed and translated in the root. Also, most of the Ribo-seq reads are mapped to the annotated reading frames in these two isoforms (**Figure 3A and 3B**). The isoform 3 is not significantly expressed in the shoot and root based on the RNA-seq coverage of the first and last exons (**Figure 3C**). The isoform 4 is not considerably expressed and translated, either (**Figure 3D**), because there are very few Ribo-seq reads in the last exon of the isoform 4 and the Ribo-seq reads in the second to the last exon are not in the expected reading frame (instead of being red, these reads are green). This indicates that the predicted coding region of the isoform 4 is not used. In other words, if the isoform 4 is actively translated, the Ribo-seq reads in the second to the last exon should be in red, and there should be plenty of reads in the CDS region within the last exon of isoform 4. Finally, from the RNA-seq coverage, it appears that the root sample expresses an isoform with the last intron retained (**Figure 3A**), suggesting intron retention or another undefined isoform exists in the root.

**Figure 3.**
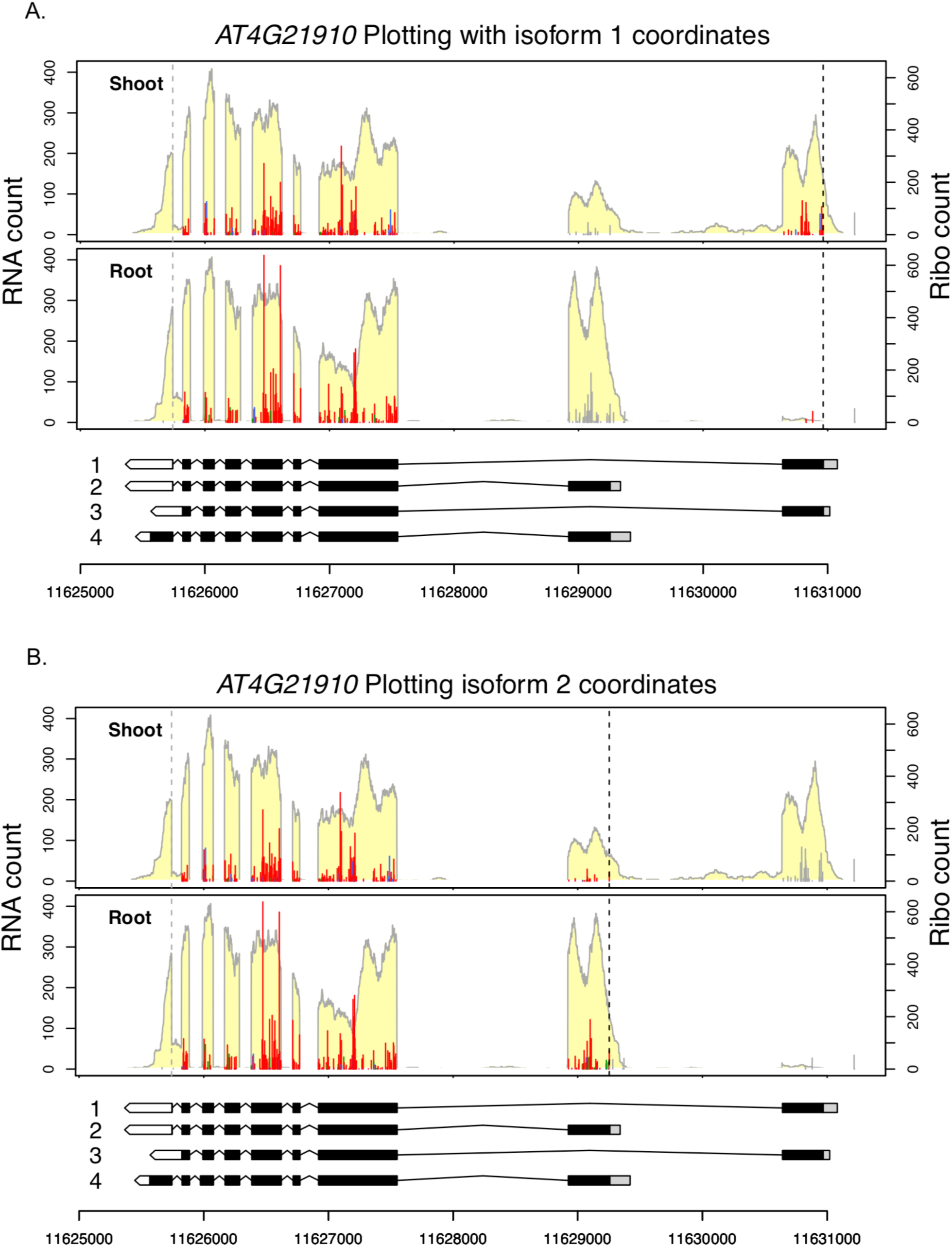
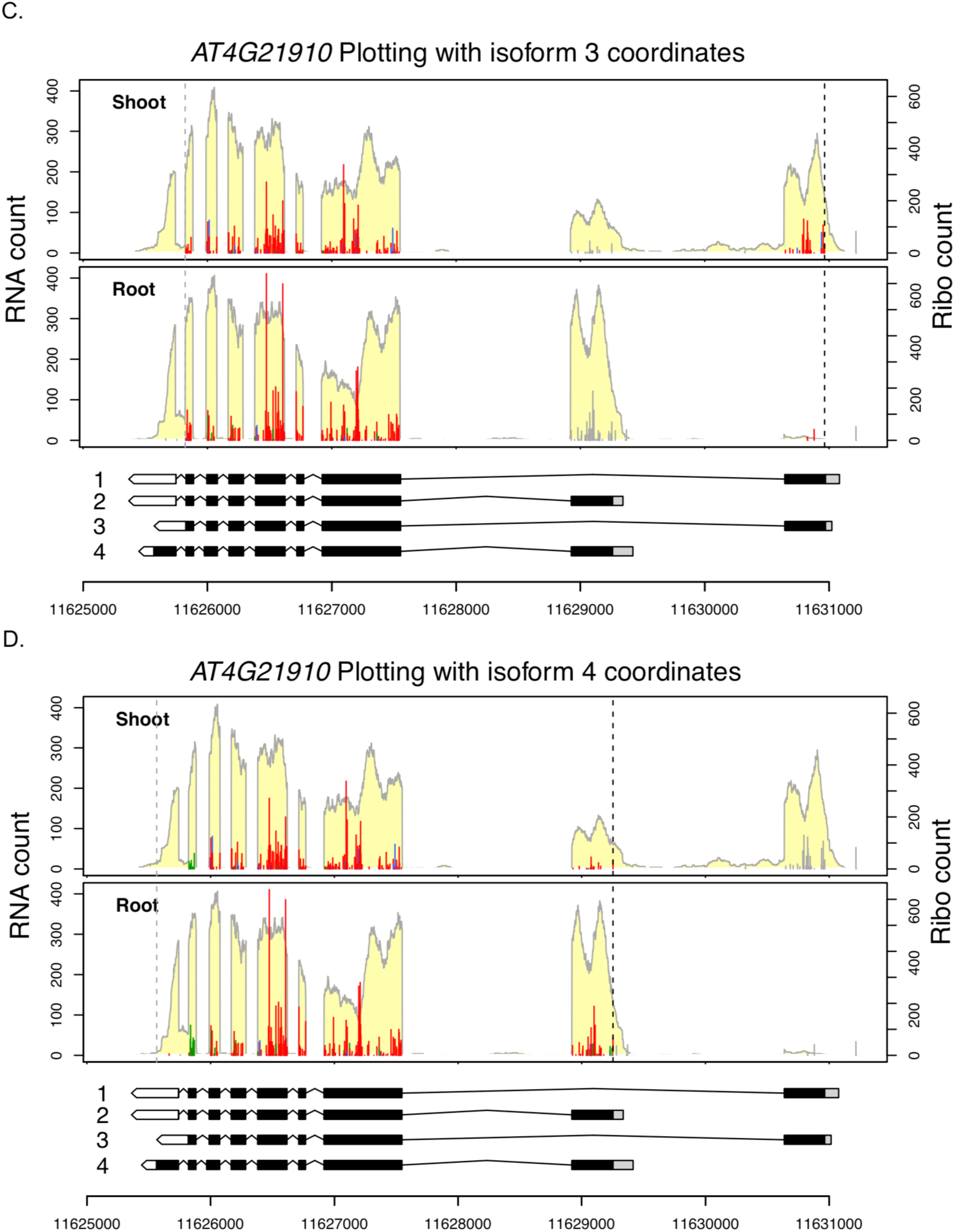
An example of plotting different transcript isoforms. The *AT4G21910* gene is plotted using PLOTc2 function with RiboPlotR comparing the Arabidopsis shoot and root. Transcript isoforms 1 to 4 of the *AT4G21910* gene are plotted in A to D, respectively. Data presentation are the same as described in the Figure 1 legend; genomic coordinates are shown below the gene models.

### Other considerations

Notice that the reads plotted in the two conditions (i.e., Root and Shoot) are not normalized. If the gene expression in the two conditions is comparable, users can normalize the reads by randomly selecting a fixed total amount of reads from the RNA-seq bam and Ribo-seq P-sites between the two conditions for plotting. However, normalizing to the same RNA-seq and Ribo-seq read counts could be easily skewed by the expression changes of highly expressed genes. For differential expression/translation analyses, users should use other software packages designated for the purposes (e.g., DESeq2 or Xtail). The RiboPlotR package is primarily used for visualizing the read distribution.

Ribo-seq is particularly useful to discover novel translational events, such as uORFs in the 5’ UTRs and small ORFs translated within presumed non-coding RNAs. Since these novel translation events are not a part of formal annotations, users have to generate a customized gtf/gff3 file to include coordinates of these novel ORFs for data visualization in RiboPlotR. Similarly, if users wish to visualize the usage of non-AUG start sites, they need to modify the CDS ranges of genes of interests in the input annotation file. More examples using RiboPlotR style to explore unknown translation events could be found in our recent publication in tomato translatome [9].

In conclusion, RiboPlotR combines a standard RNA-seq bam file, a transcriptome annotation file, and Ribo-seq P-site position file to plot the RNA-seq coverage and the Ribo-seq P-site positions in each isoform considered. It could be used to investigate the translation of specific transcript isoforms, uORFs, and other novel translational events.

## References

1. Ingolia NT, Ghaemmaghami S, Newman JRS, Weissman JS (2009) Genome-Wide Analysis in Vivo of Translation with Nucleotide Resolution Using Ribosome Profiling. Science (80-) 324:218–223

2. Kiniry SJ, O’Connor PBF, Michel AM, Baranov P V (2019) Trips-Viz: a transcriptome browser for exploring Ribo-Seq data. Nucleic Acids Res 47:D847–D852

3. Lawrence M, Huber W, Pagès H, Aboyoun P, Carlson M, Gentleman R, Morgan MT, Carey VJ (2013) Software for Computing and Annotating Genomic Ranges. PLoS Comput Biol 9:e1003118

4. Morgan M, Pagès H, Obenchain V HN (2019) Rsamtools: Binary alignment (BAM), FASTA, variant call (BCF), and tabix file import. R package version 2.0.0. http://bioconductor.org/packages/Rsamtools

5. Lawrence M, Gentleman R, Carey V (2009) rtracklayer: an R package for interfacing with genome browsers. Bioinformatics 25:1841–1842

6. Li H, Handsaker B, Wysoker A, Fennell T, Ruan J, Homer N, Marth G, Abecasis G, Durbin R (2009) The Sequence Alignment/Map format and SAMtools. Bioinformatics 25:2078–2079

7. Dobin A, Davis CA, Schlesinger F, Drenkow J, Zaleski C, Jha S, Batut P, Chaisson M, Gingeras TR (2013) STAR: ultrafast universal RNA-seq aligner. Bioinformatics 29:15–21

8. Hsu PY, Calviello L, Wu H-YL, Li F-W, Rothfels CJ, Ohler U, Benfey PN (2016) Super-resolution ribosome profiling reveals unannotated translation events in Arabidopsis. Proc Natl Acad Sci U S A 113:E7126–E7135

9. Wu H-YL, Song G, Walley JW, Hsu PY (2019) The tomato translational landscape revealed by transcriptome assembly and ribosome profiling. Plant Physiol pp.00541.2019

